# Interaction of *Plasmodium falciparum* Casein kinase 1 (PfCK1) with components of host cell protein trafficking machinery

**DOI:** 10.1101/617571

**Authors:** Mitchell B Batty, Ralf B Schittenhelm, Christian Doerig, Jose Garcia-Bustos

## Abstract

During infection, the *Plasmodium falciparum* casein kinase 1 (PfCK1) is secreted to the extracellular medium and appears on the RBC membrane during trophozoite stage of development. We attempted to identify a mechanism that describes the secretion of PfCK1 and its appearance on the RBC membrane and suspected a mechanism involving multiple host proteins may be utilised. Indeed, we found that the host proteins GTPase-activating protein and Vps9 domain-containing protein (GAPVD1) and Sorting nexin 22 (SNX22), which have described functions in membrane trafficking in higher eukaryotes, consistently co-purify with PfCK1 suggesting the parasite utilises trafficking pathways previously thought to be inactive in RBCs. Further, reciprocal immunoprecipitation experiments with GAPVD1 identified parasite proteins suggestive of a recycling pathway hitherto only described in higher eukaryotes to recycle membrane proteins. Thus, we have identified components of a trafficking pathway involving parasite proteins that act in concert with host proteins which we hypothesise coordinate the trafficking of PfCK1 during infection.

## Introduction

Post-translational modifications (PTMs) are essential biochemical switches that regulate protein interactions and thereby many cellular functions. A PTM of major importance is protein phosphorylation, a dynamic process regulated by protein kinases and phosphatases, with over 500 protein kinase and more than 200 protein phosphatase coding genes identified in humans [1, 2]. Dysregulation of this essential modification contributes to the establishment of diseases such as cancer [3]. In the malaria parasite *Plasmodium falciparum*, the number of kinases is smaller, with 86 annotated protein kinases though this number varies depending on the parameters used in each study [4-7]. Of these, 36 have been identified as likely essential for asexual blood stage development [8]] – the stages of development responsible for disease pathology and the target of most current antimalarials. In view of the success in targeting protein kinases in diseases like cancer [9], *P. falciparum* protein kinases are viewed as potential targets for drugs with novel modes of action and therefore free of pre-existing resistance mechanisms.

Casein kinase 1 (CK1) is a highly conserved eukaryotic PK with seven isoforms identified in mammalian cells, some of which are expressed as multiple splice variants [10]. CK1 activity is important for normal cellular function and is involved in many biological processes including Wnt/β-catenin signalling [11], regulation of the cell cycle [12] and endocytosis [13]. Mutations in CK1 or inhibition of its kinase activity results in the dysregulation of biochemical pathways and can lead to the development of disease including cancer [14]; CK1 is therefore an attractive therapeutic target. CK1 recognises and phosphorylates the canonical sequence *pS/pT X*_*1-2*_ *<u>S/T</u>* where pS/pT denotes a “primed” phosphorylated Ser/Thr residue and X is any amino acid; phosphorylation occurs on the next Ser/Thr residue (underlined)[15]. Early investigations into the determinants of substrate recognition by CK1 also showed that a cluster of 3-5 acidic residues upstream of a Ser/Thr is capable of priming substrates for phosphorylation [16]. In *P. falciparum*, CK1 is present as a single 36kDa isozyme and is one of the 36 protein kinases identified as likely essential for asexual blood stage development. Recent work has identified the cellular distribution and possible processes that involve PfCK1 [17]. During the ring to early trophozoite phase, a large proportion of PfCK1 appears to be associated with the host red blood cell membrane and may be an ectokinase (a protein kinase with an extracellularly facing kinase domain). PfCK1 activity has also been detected in the supernatant of trophozoite stage parasite cultures, indicating that this protein is also secreted during the asexual lifecycle [17]. These observations are similar to those made in *Leishmania*, where ectokinase status and secretion to the culture medium have been observed for one of the CK1 homologues present in this species [18, 19]. Progression to schizonts shows PfCK1 becoming less associated with the membrane and more localised to the parasite itself, finally appearing in merozoites only. This membrane association is interesting because PfCK1 has no PEXEL (<u>P</u>lasmodium <u>EX</u>port <u>EL</u>ement) motif or any form of recognisable export signal sequence [20, 21], meaning it is probably not subject to proteolytic processing in the endoplasmic reticulum (ER) and subsequently translocated across the parasitophorous vacuole membrane (PVM) to the host cytoplasm via the PTEX transposon [21, 22]. Furthermore, there appears to be no interaction of PfCK1 with Maurer’s clefts either. These are parasite-derived membranous structures associated with the RBC cytoskeleton able to transport parasite proteins to the RBC membrane [23]. This indicates that trafficking and secretion of PfCK1 may occur via a hitherto undescribed mechanism. We hypothesised that trafficking of PfCK1 to the host cell plasma membrane involves trafficking machinery and probably interactions with host proteins. In order to identify such proteins, we used affinity purification and mass spectrometry to analyse co-precipitates obtained from transgenic parasites expressing PfCK1-GFP from the endogenous locus and verified candidate interactors by reciprocal co-immunoprecipitation. To test this hypothesis, we set out to determine whether host erythrocytes protein associate with PfCK1, using immunoprecipitation of PfCK1 from infected red blood cells lysates coupled to mass spectrometry analysis. This method had been successfully used previously to document the parasite interactome of PfCK1 using purified parasites as the starting material [17]. Briefly, we prepared extracts from synchronised red blood cell cultures infected with a 3D7 parasite line expressing GFP-tagged PfCK1from its endogenous locus and magnet-purified infected erythrocytes at trophozoite stages by. Lysates were prepared by incubating whole blood cultures (equivalent to 1×10^8^ infected red blood cells) with 2mL of filter sterile lysis buffer (10mM Tris, pH 7.5, 150mM NaCl, 10mM NaF, 10mM β-glycerophosphate, 0.1mM Na_3_VO_4_, 1mM PMSF, 1X Protease inhibitor cocktail, 0.5% IGEPAL, 0.5mM EDTA) for 30mins at 4°C with constant rotation and subjected to immunoprecipitation using GFP Trap® beads. Beads were washed three times with 1mL of wash buffer (10mM Tris, pH 7.5, 500mM NaCl, 0.5mM EDTA, 10mM NaF, 10mM β-glycerophosphate, 0.1mM Na_3_VO_4_, 1mM PMSF, 1% IGEPAL and 1X Protease inhibitor cocktail) and eluted in 50μl of 2x SDS sample buffer. Protein eluates were briefly run into a 4-12% SDS-PAGE gel (∼5mins) and peptides were generated by in-gel tryptic digestion and analysed by mass spectrometry.

A 3D7 wild-type parasite line expressing its native (untagged) PfCK1 was used as a negative control. The raw spectral data from PfCK1-GFP and wild-type PfCK1 experiments was subjected to principal component analysis (PCA) (supplementary figure 1) and used to interrogate both the human and *Plasmodium* proteomes. Subsequent statistical analysis was performed in MaxQuant to identify high-probability interactors and are presented in a volcano plot. A good separation between PfCK1-GFP and wild-type samples was observed in the PCA (supplementary figure S1) and PfCK1 was identified in precipitates obtained from PfCK1-GFP samples but not from wild-type-derived samples, as expected (Figure 1A). We reproducibly identified ten proteins as significantly different between PfCK1-GFP and wild-type samples (supplementary table 1), the strongest hits being the host proteins GTPase-activating protein-VPS9 domain-containing protein 1 (GAPVD1), sorting nexin 22 (SNX22) and human CK1α. This is consistent with published observations that GAPVD1 and SNX24 (which clusters with SNX22 in phylogenetic trees) co-purify with the mammalian CK1 isoform CK1ε in bait-prey pull-down experiments [24, 25]. Sorting nexins anchor cargo proteins to membranes enriched in phosphatidylinositol-3-phosphate (PtdIns3P) [26] *via* their Phox homology (PX) domain (Figure 1C) and have been implicated in protein secretion [27]. Whilst there are no functions described for SNX22, other members of this family, for instance SNX3, are well-studied and known to function in directing retromer-mediated vesicle transport to the *trans*-Golgi network (TGN), a conserved trafficking pathway first identified in yeast which guides the retrieval, sorting, recycling and retrograde transport of membrane receptors [28]. Furthermore, phosphorylation within the PX domain of SNX3 (Figure 1C) abolishes PtdIns binding, thus resulting in cytosolic localisation and demonstrates the modulation of SNX function by protein kinases [29]. GAPVD1 contains an evolutionarily conserved Vps9 domain (Figure 1C) that functions as a guanine nucleotide exchange factor (GEF) for Ras-related GTP-binding proteins [30] and has established functions in receptor-membrane transport [31, 32]. Trafficking of GAPVD1 to the plasma membrane also results in the synthesis and turnover of PtdIns3 by coordinating the GTPase Rab5 and PtdIns3P-kinases (PI3K) [33]. *P. falciparum* does not encode homologs to these host proteins but does encode Rab GTPases such as PfRab5 (involved in trafficking PfCK1 to the parasite membrane [34]) and PfRab7 which co-localises with components of the retrograde transport complex in punctate structures within close proximity to the Golgi apparatus ([35] and reviewed in [36]). In light of their various roles in protein trafficking and their high abundance PfCK1-GFP immunoprecipitates, we experimentally tested the association of PfCK1 with GAPVD1 (UniProt ID: Q14C86) and SNX22 (UniProt ID: Q96L94). For these two host proteins, we performed Western blot analysis of immunoprecipitates obtained with GFP Trap beads, probing each blot with cognate antibodies. In both cases, presence of the target protein was confirmed in precipitates obtained from parasite samples expressing GFP-tagged PfCK1, but not in samples expressing the wild-type kinase (Figure 1B). Several bands were observed in blots probed with anti-GAPVD1 antibody, consistent with indicating multiple phosphorylation sites. Indeed, eight phosphopeptides that mapped to various regions of GAPVD1 (Table 1) were identified in PfCK1-GFP precipitates but not WT samples, with pS903 and pS1104 phosphosites present in all three PfCK1-GFP replicates. Two of the eight identified phosphosites (pS1104 and pS972) are located downstream of a cluster of Asp residues, consistent with the ability of CK1 proteins to recognise acidic sequences upstream of a target Ser/Thr residue to prime phosphorylation [16]. None of these phosphopeptides map to the Vps9 domain (guanine nucleotide exchange factor) or the Ras-GTPase activating protein (Ras-GAP) domain (Figure 1C), suggesting they probably do not regulate catalysis but likely function in regulating other processes, such as adaptor protein interactions. We next verified the interaction between GAPVD1 and PfCK1 in infected cells by performing the reciprocal immunoprecipitation using an antibody against GAPVD1. Wild-type 3D7 parasites were harvested at trophozoite stages by magnet purification, lysed as described above and incubated with anti-GAPVD1 antibody bound to Protein A sepharose overnight at 4°C. Uninfected RBCs were used as a control. Proteins were eluted and peptides generated by in-gel tryptic digestion and analysed by mass spectrometry, checking spectra against human and *Plasmodium* proteomes as previously outlined. Statistical analysis was performed in MaxQuant and the results from independent triplicate experiments are displayed as a volcano plot. Good separation between wild-type infected (iRBCs) and uninfected (uRBCs) red blood cells was observed in the PCA (supplementary Figure 2) and PfCK1 was detected in the subset of 48 proteins identified as high probability GAPVD1 interactors (Figure 2A), providing independent confirmation that PfCK1 forms a protein complex with GAPVD1. Interestingly, we identified the *P. falciparum* Vacuolar protein sorting-associated protein 51 (PfVps51) as a high probability interactor in our reciprocal GAPVD1 data set (Figure 2A). In eukaryotes Vps51 forms a core component of the Golgi-associated retrograde protein (GARP) tethering complex that regulates endosome fusion with the *trans*-golgi network (TGN), mediated by interactions with SNARE proteins [37]. As with SNX proteins, GARP has also been implicated in retromer-dependent protein secretion [37]. Further, the SNARE associated protein snapin is a substrate of CK1δ and localises to membranes of the Golgi apparatus [38] lending support to the possible involvement of PfVps51 in the trafficking of PfCK1. Importantly, Vps51 and SNARE proteins are present in *Leishmania a*nd *Plasmodium* [39, 40] indicating this pathway is conserved in parasitic protozoa, consistent with the identification of protein trafficking and export being as of the most represented functional groups in our pathway analysis of parasite proteins identified in GAPVD1 precipitates (Figure 2B).

**Table 1.**
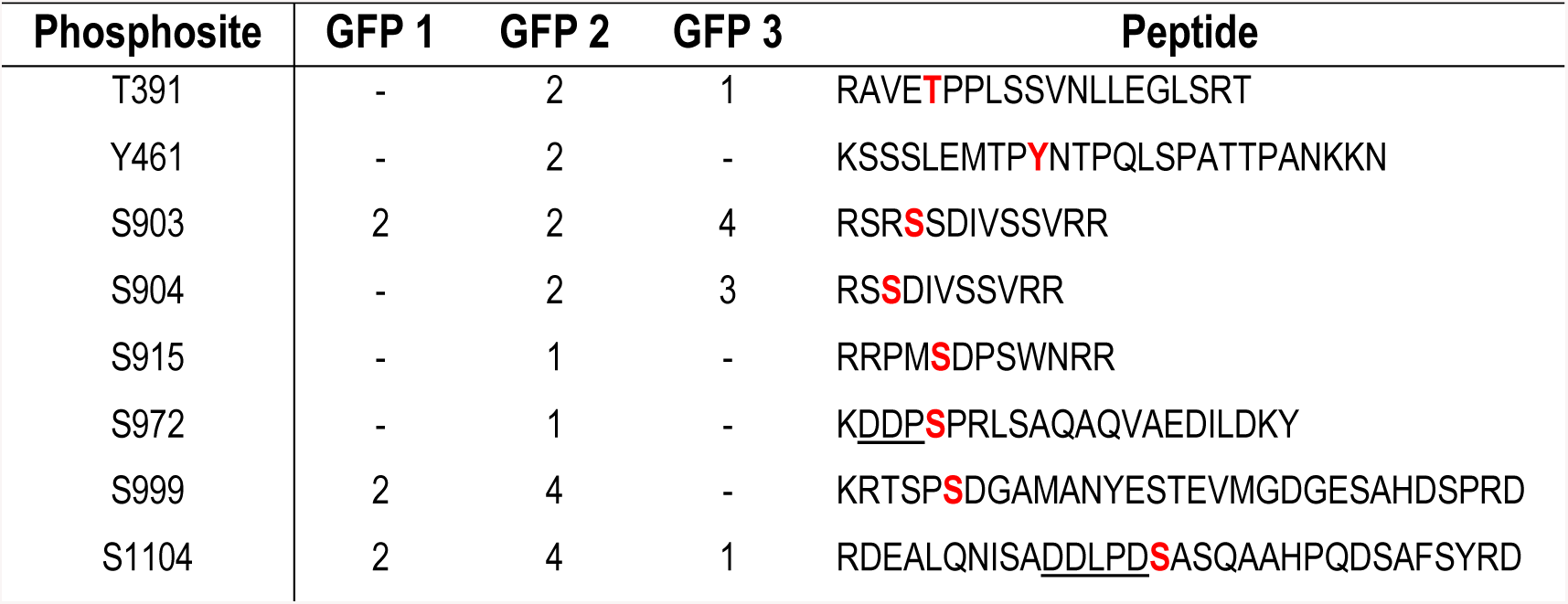
Number of GAPVD1 phosphopeptides obtained from PfCK1-GFP precipitations. Bold and red indicates the identified phosphosites and underlined are possible consensus sequence for CK1 phosphorylation.

**Figure 1.**
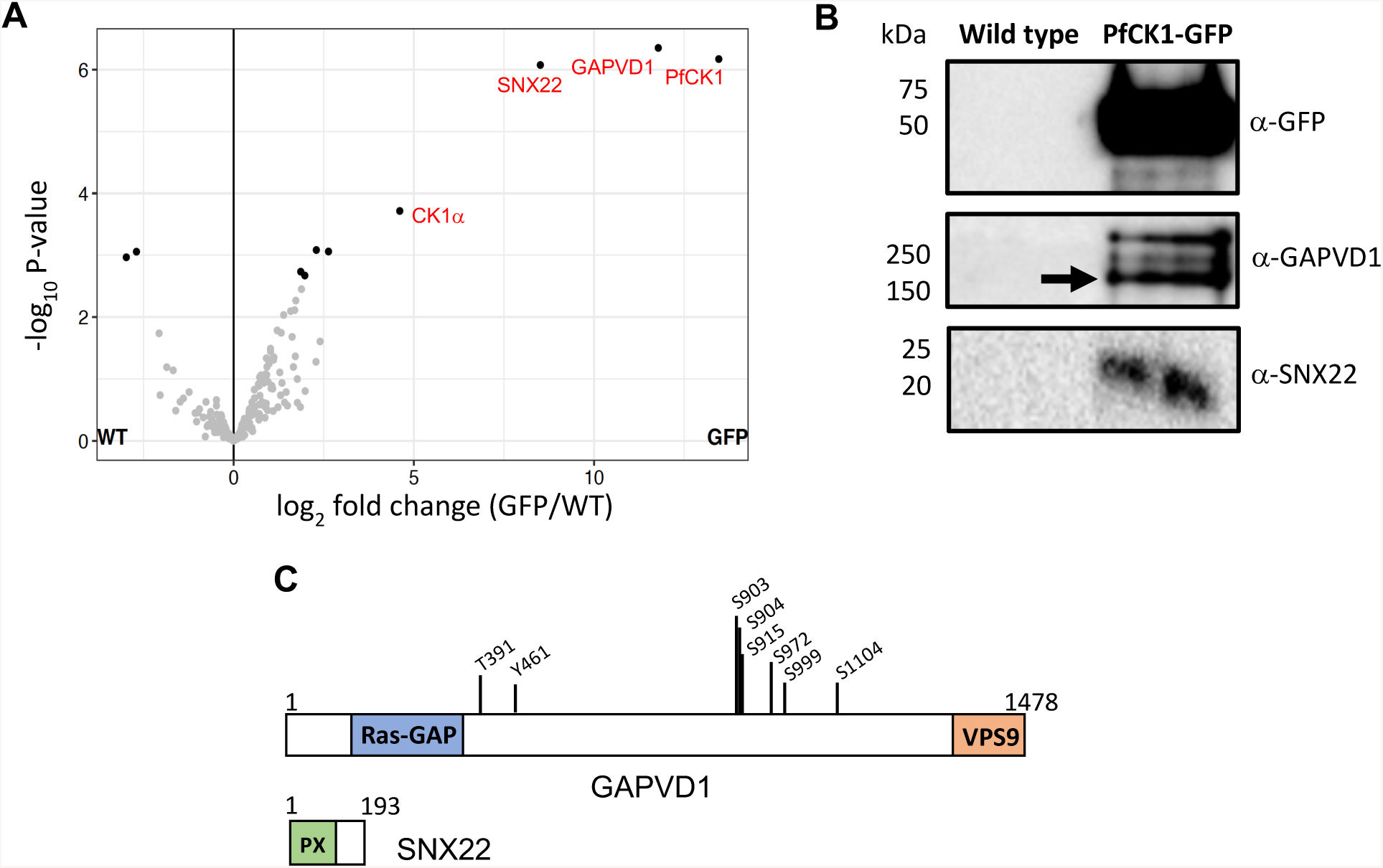
Label-free quantitative analysis of human interactors of PfCK1. Whole blood lysates were prepared from a 3D7 parasite line expressing PfCK1-GFP from its endogenous locus and harvested at trophozoite stages by magnet purification. Wild-type 3D7 parasites expressing the native untagged PfCK1 were used as a negative control. Lysates were incubated with GFP Trap® beads on an end-over-end rotator for 2hrs at 4°C. Beads were washed 3x with 1mL wash buffer and eluted in 50μl 2x SDS sample buffer (Biorad). Peptides were generated by in-gel tryptic digestion and analysed by mass spectrometry. The raw data files were analyzed using MaxQuant to obtain protein identifications and their respective label-free quantification values using in-house standard parameters. Data were normalized based on the assumption that the majority of proteins do not change between the different conditions. (A) Volcano plot representing the logarithmic ratio of LFQ values between PfCK1-GFP/WT samples, plotted against the negative logarithm of p-values obtained from Student’s t-test of triplicate experiments (FDR = 0.1). High probability interactors (solid black circles) were determined with an adjusted p-value cut-off of ≤0.05 and log fold change ≥1.5. (B) Analysis of immunoprecipitation experiments by Western blot probed with rabbit antibodies against GFP, GAPVD1 and SNX22. Arrow indicates the estimated size of GAPVD1. (C) Schematic diagram of the domain organisation of GAPVD1 and SNX22 with each functional domain denoted; GTPase activating protein (Ras-GAP), Guanine nucleotide exchange factor (VPS9) and Phox homology domain (PX). Location of GAPVD1 phosphosites identified in PfCK1-GFP precipitates are annotated.

**Figure 2.**
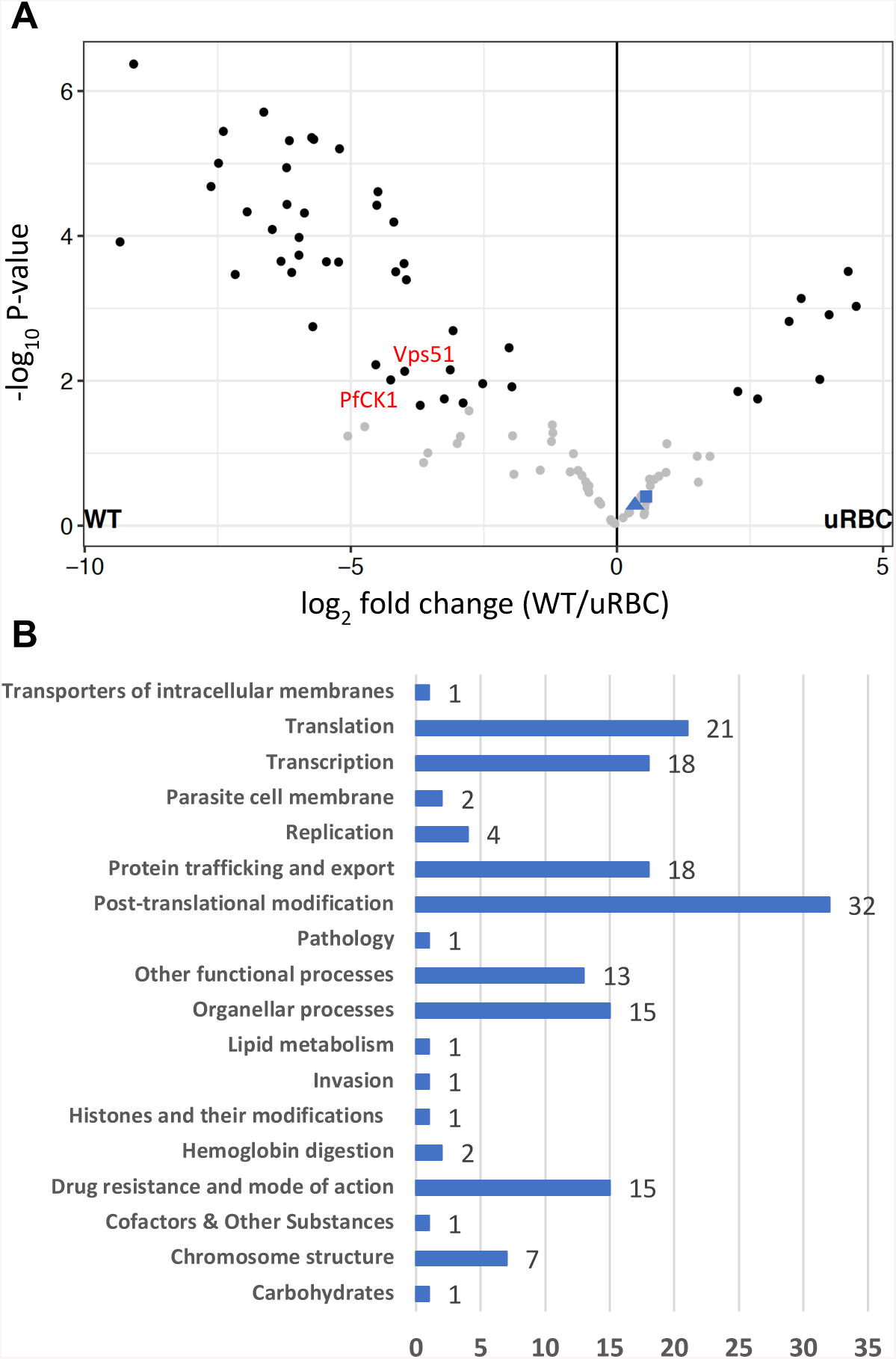
Label free quantitative analysis of parasite interactors of GAPVD1. Protein lysates were prepared from wild-type 3D7 parasites harvested at trophozoite stages by magnet purification. Lysates were incubated with 2μg of GAPDV1 antibody (Abcam) bound to Protein A sepharose beads (Sigma-Aldrich) washed in lysate buffer on an end-over-end rotator overnight at 4°C. Beads were washed with 0.5M NaCl and 1% v/v IGEPAL and eluted in 2x SDS sample buffer (Biorad). As outlined previously, generated peptides were analysed by mass spectrometry the raw data files were analyzed using MaxQuant to obtain protein identifications and their respective label-free quantification values using in-house standard parameters. (A) Volcano plot representing the logarithmic ratio of LFQ values between iRBS/uRBCs incubated with GAPDV1 antibody which are plotted against the negative logarithmic p-values obtained from student’s t-test of triplicate experiments. Cut-off values of ≤0.05 for p-value and ≥1.5 log fold change was used to separate high probability interactors (black) from background (grey) proteins. Key parasite proteins are annotated in red and GAPVD1 and SNX22 (solid triangle and square respectively) are also marked. (B) Parasite proteins identified as high-probability interactors of GAPVD1 distributed across the various metabolic pathways of the malaria parasite. The histogram represents the number of parasite proteins mapped to each pathway described in the Metabolic Pathways of Malaria Parasites website (http://mpmp.huji.ac.il//).

In summary, we provide evidence for (i) the presence of GAPVD1 and SNX22-dependent protein trafficking pathways in human RBCs, and (ii) the interaction of PfCK1 with these host erythrocyte proteins. Furthermore, our co-precipitation data suggests that both GAPVD1 and SNX22 function in a complex, providing evidence of interplay between pathways. Their exact functions in RBCs still remain unknown however we are currently exploring human interactors of SNX22 and GAPVD1 in an attempt to gain a further insight into their possible roles in RBCs. Nevertheless, the identification of PfCK1 in this complex suggests that during infection, *P. falciparum* parasites commandeer remanent components of the endosomal trafficking machinery in RBCs, repurposing these for trafficking parasite proteins to membranes and possibly also to the external medium. Our data also suggests that *P. falciparum* parasites contain components of an endosomal recycling pathway that may provide a link between parasite endosomal processes and host trafficking machinery.

