## Supplemental Tables and Figures for "Interaction of *Plasmodium falciparum* Casein kinase 1 (PfCK1) with components of host cell protein trafficking machinery"

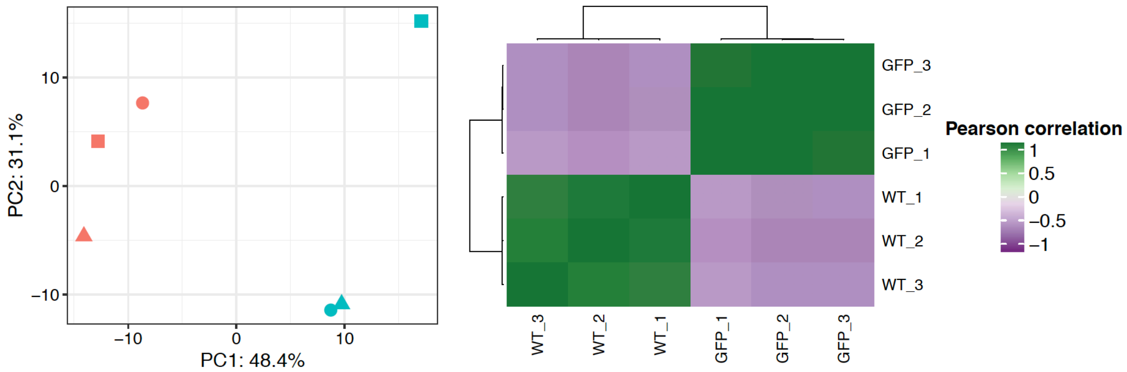


**Supplementary Figure S1.** Exploratory analysis of replicate data sets obtained from PfCK1-GFP immunoprecipitates. (A) Principal component analysis shows good separation between wild-type (blue) and PfCK1-GFP (red) samples (B) Pearson correlation matrix showing linear relationship between samples and conditions. A strong correlation is indicated in green (value of 1) and weak as purple (value of -1).

**Supplementary table S1.** Proteins identified as high probability interactors in PfCK1-GFP immunoprecipitates. Hits are ranked highest to lowest by -log Student's t-test (p-value)

| Protein names | Protein ID | Gene | -log Student's t-test p-value | $log_2$ fold change |
|---|---|---|---|---|
| GTPase-activating protein and VPS9 domain-containing protein 1 | Q14C86 | GAPVD1 | 4.126 | 11.784 |
| Sorting nexin-22 | Q96L94 | SNX22 | 4.125 | 7.410 |
| 14-3-3 protein epsilon | P62258 | YWHAE | 2.930 | 2.294 |
| Casein kinase I isoform alpha | P48729 | CSNK1A1 | 2.664 | 4.978 |
| Probable ubiquitin carboxyl-terminal hydrolase FAF-X | Q93008 | USP9X | 2.574 | 1.975 |
| T-complex protein 1 subunit epsilon | E7ENZ3 | CCT5 | 2.275 | 1.882 |
| Heat shock 70 kDa protein | P0DMV9 | HSPA1A | 2.156 | 1.725 |
| T-complex protein 1 subunit zeta | P40227 | CCT6A | 2.050 | 1.582 |
| Protein S100-A9 | P06702 | S100A9 | 1.980 | -3.053 |
| Desmocollin-1 | Q08554 | DSC1 | 1.978 | -2.646 |
| 14-3-3 protein zeta/delta | P63104 | YWHAZ | 1.922 | 1.693 |
| T-complex protein 1 subunit theta | P50990 | CCT8 | 1.380 | 1.623 |
| T-complex protein 1 subunit alpha | P17987 | TCP1 | 1.348 | 1.116 |
| Casein kinase I | Q8IHZ9 | PfCK1 | 3.914 | 13.463 |
| 40S ribosomal protein SA | Q8IJD4 | PF10_0264 | 3.070 | 1.862 |
| Heat shock protein 70 | Q8IB24 | PF08_0054 | 2.498 | 2.632 |
| Heat shock protein 71 | Q8I2X4 | PFI0875w | 2.415 | 1.387 |
| Tubulin beta chain | Q7KQL5 | PF10_0084 | 1.800 | 1.319 |

*Proteins identified as significant based on defined cut-off values of ≤1.3 for -log student's t-test (p-value equivalent to ≤0.05) and a value of ≥1 for $log_2$ fold change (≥2-fold change). Data is ranked highest to lowest according to p-value. Highlighted in green are parasite proteins identified as significant in PfCK1-GFP immunoprecipitates.*

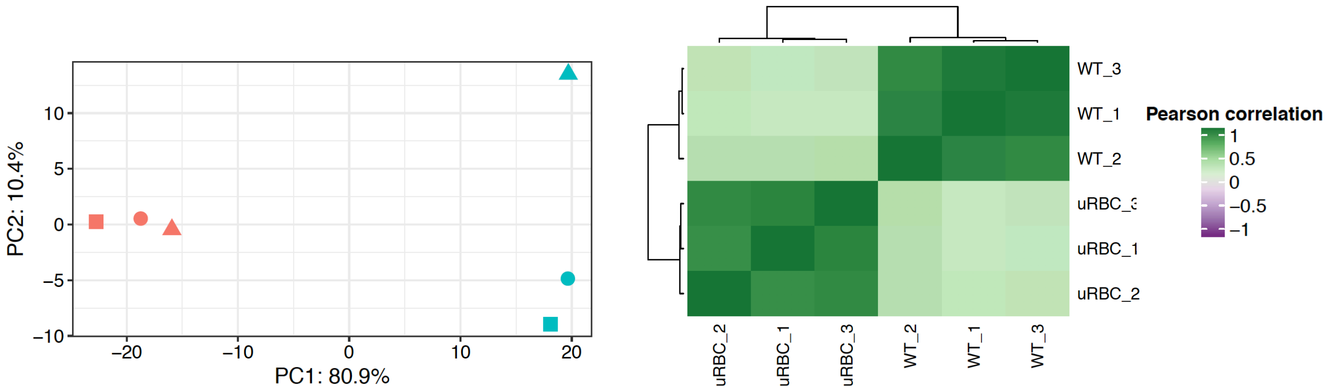


**Supplementary Figure S2.** Exploratory analysis of replicate data sets obtained from GAPVD1 immunoprecipitates. (A) Principal component analysis shows good separation between wild-type infected (blue) and uninfected (red) red blood cells. (B) Pearson correlation matrix showing linear relationship between samples and conditions. A strong correlation is indicated in green (value of 1) and weak as purple (value of -1).

**Supplementary table S2.** Proteins identified as high probability interactors in GAPVD1 immunoprecipitates from RBCs infect with wild-type parasites and uRBCs used as control samples. Hits are ranked highest to lowest by -log Student's t-test (p-value).

| Protein names | Protein ID | Gene names | -log Student's t-test (P-value) | Log2 fold change |
|---|---|---|---|---|
| DNA/RNA-binding protein Alba3 | Q8IJX8 | PF10_0063 | 5.262 | 5.157 |
| Polyadenylate-binding protein | Q8I5H4 | PFL1170w | 4.984 | 6.274 |
| Heat shock protein 70 | Q8II24 | PF11_0351 | 4.949 | 4.737 |
| Heat shock protein 70 | Q8I2X4 | PFI0875w | 4.222 | 6.154 |
| GTP binding nuclear protein | Q7KQK6 | PF11_0183 | 4.219 | 5.361 |
| Eukaryotic initiation factor 4A | Q8IKF0 | H45 | 3.959 | 5.932 |
| Heat shock protein 70 | Q8IB24 | PF08_0054 | 3.945 | 7.394 |
| Actin-1 | Q8I4X0 | PFL2215w | 3.743 | 6.078 |
| Heat shock protein 90 | Q8IC05 | PF07_0029 | 3.741 | 9.554 |
| 60S acidic ribosomal protein P2 | O00806 | MAL3P3.19 | 3.679 | 5.626 |
| S-adenosylmethionine synthase | Q7K6A4 | PfSAMS | 3.589 | 4.225 |
| 60S ribosomal protein L28 | Q8IHU0 | PF11_0437 | 3.483 | 3.922 |
| Heat shock protein 110 | Q8IC01 | PF07_0033 | 3.297 | 8.137 |
| 14-3-3 protein | C0H4V6 | MAL8P1.69 | 3.246 | 6.021 |
| 60S acidic ribosomal protein P0 | Q8II61 | PF11_0313 | 3.246 | 6.569 |
| Karyopherin beta | Q8I3M5 | PFE1195w | 3.206 | 4.369 |
| Uncharacterised protein | Q8IKG9 | PF14_0636 | 3.014 | 6.983 |
| Glyceraldehyde-3-phosphate dehydrogenase | Q8IKK7 | GAPDH | 3.004 | 5.807 |
| Proline--tRNA ligase | Q8I5R7 | proRS | 2.955 | 4.985 |
| Glycophorin-binding protein | Q8I6U8 | GBP | 2.93 | 6.247 |
| 40S ribosomal protein S12 | O97249 | RPS12 | 2.881 | 5.131 |
| NLI interacting factor-like phosphatase, putative | Q8IJR8 | PF10_0124 | 2.77 | 8.109 |
| Vacuolar protein sorting-associated protein 51, putative | Q8I2A9 | PFA_0155c | 2.729 | 3.921 |
| Casein kinase I | Q8IHZ9 | PfCK1 | 2.559 | 4.422 |
| Plasmepsin II | Q8I6V3 | PF14_0077 | 2.323 | 3.487 |
| 60S ribosomal protein L5 | Q8ILL3 | PF14_0230 | 2.285 | 3.547 |
| Ornithine aminotransferase | Q6LFH8 | OAT | 2.24 | 3.826 |

| | | | | |
|---|---|---|---|---|
| 40S ribosomal protein S25 | Q8ILN8 | PF14_0205 | 2.228 | 2.192 |
| V-type H(+)-translocating pyrophosphatase, putative | Q8IKR1 | PF14_0541 | 2.168 | 3.499 |
| Elongation factor 1-alpha | Q8I0P6 | PF3D7_1357000 | 2.11 | 7.661 |
| HAP protein | Q8IM15 | PF14_0078 | 2.013 | 3.989 |
| 40S ribosomal protein S19 | Q8IFP2 | PFD1055w | 1.92 | 2.294 |
| Small exported membrane protein 1 | Q8IC43 | PF07_0007 | 1.568 | 3.82 |
| Spectrin beta chain, erythrocytic | Q01082 | SPTB | 2.54 | -4.348 |
| Protein-L-isoaspartate(D-aspartate) O-methyltransferase | P22061 | PCMT1 | 2.472 | 2.024 |
| Casein kinase I isoform alpha | P48729 | CSNK1A1 | 2.459 | -4.317 |
| Protein 4.1 | Q9H4G0 | EPB41 | 2.39 | -3.465 |
| Spectrin alpha chain, erythrocytic 1 | P02549 | SPTA1 | 2.123 | -3.237 |
| Heat shock 70 kDa protein 6 | P17066 | HSPA6 | 1.928 | 5.501 |
| Erythrocyte band 7 integral membrane protein | P27105 | STOM | 1.556 | 0.815 |
| Heat shock cognate 71 kDa protein | P11142 | HSPA8 | 1.528 | -0.941 |
| Flavin reductase (NADPH) | P30043 | BLVRB | 1.525 | 1.212 |
| Tropomodulin-1 | P28289 | TMOD1 | 1.46 | -2.805 |
| Ankyrin-1 | Q01484 | ANK1 | 1.42 | -2.274 |
| Dematin | Q08495 | DMTN | 1.369 | -2.876 |
| Tropomyosin alpha-3 chain | Q3SX28 | TPM3 | 1.324 | -2.894 |
| Sorting nexin-22 | Q96L94 | SNX22 | 0.94 | 0.727 |
| GTPase-activating protein and VPS9 domain-containing protein 1 | Q14C86 | GAPVD1 | 0.822 | 0.585 |

*Proteins identified as significant based on defined cut-off values of ≤1.3 for -log student's t-test (p-value equivalent to ≤0.05) and a value of ≥1 for $log_2$ fold change (≥2-fold change). Data is ranked highest to lowest according to p-value. Highlighted in blue are host RBC proteins identified as significant in GAPVD1 immunoprecipitates.*
